# An Orientationally Averaged Version of the Rotne-Prager-Yamakawa Tensor Provides A Fast But Still Accurate Treatment Of Hydrodynamic Interactions In Brownian Dynamics Simulations Of Biological Macromolecules

**DOI:** 10.1101/2023.04.21.537865

**Authors:** John W. Tworek, Adrian H. Elcock

## Abstract

The Brownian dynamics (BD) simulation technique is widely used to model the diffusive and conformational dynamics of complex systems comprising biological macromolecules. For the diffusive properties of macromolecules to be described correctly by BD simulations, it is necessary to include hydrodynamic interactions (HI). When modeled at the Rotne-Prager-Yamakawa (RPY) level of theory, for example, the translational and rotational diffusion coefficients of isolated macromolecules can be accurately reproduced; when HIs are neglected, however, diffusion coefficients can be underestimated by an order of magnitude or more. The principal drawback to the inclusion of HIs in BD simulations is their computational expense, and several previous studies have sought to accelerate their modeling by developing fast approximations for the calculation of the correlated random displacements. Here we explore the use of an alternative way to accelerate calculation of HIs, i.e., by replacing the full RPY tensor with an orientationally averaged (OA) version which retains the distance dependence of the HIs but averages out their orientational dependence. We seek here to determine whether such an approximation can be justified in application to the modeling of typical proteins and RNAs. We show that the use of an OA RPY tensor allows translational diffusion of macromolecules to be modeled with very high accuracy at the cost of rotational diffusion being underestimated by ∼25%. We show that this finding is independent of the type of macromolecule simulated and the level of structural resolution employed in the models. We also show, however, that these results are critically dependent on the inclusion of a non-zero term that describes the divergence of the diffusion tensor: when this term is omitted from simulations that use the OA RPY model, unfolded macromolecules undergo rapid collapse. Our results indicate that the orientationally averaged RPY tensor is likely to be a useful, fast approximate way of including HIs in BD simulations of intermediate-scale systems.

## Introduction

The Brownian dynamics (BD) simulation algorithm developed by Ermak and McCammon [1] is a widely used approach for simulating the conformational and diffusive dynamics of complex biomolecular systems while including the effects of hydrodynamic interactions (HIs). In most applications of the algorithm, HIs are described at the pairwise Rotne-Prager-Yamakawa (RPY) level of theory [2, 3], and it has been shown that when used in simulations of macromolecules represented by flexible, bead-spring models it provides an accurate means of modeling their translational and rotational diffusion (e.g. [4, 5]). A major drawback of the BD-HI method, however, is the significant computational expense associated with accounting for the RPY HIs. There are two principal factors that contribute to this expense (see Methods), but both can be attributed to the fact that the RPY diffusion tensor, which describes the HIs operating between all pairs of beads, has dimensions of 3N × 3N, where N is the number of beads in the system. Here we consider the possibility of accelerating BD-HI simulations by replacing the full RPY tensor with an orientationally averaged version that allows HIs to fluctuate during the simulation and allows them to retain their distance dependence, but reduces the diffusion tensor’s dimensions to N × N, thereby allowing potential simulation speedups of between 3-fold and 27-fold (see below). While there is a long history of using this orientational averaging approximation to simplify the treatment of HIs in both theory (e.g. [6]) and numerical calculations (e.g. [7]), this is, to our knowledge, the first report of its direct use in BD-HI simulations. In common with the findings of previous studies of the effects of orientational averaging [7–9], we show here that it results in relatively little loss of accuracy in the treatment of translational diffusion but leads to an ∼25% underestimation of the rate of rotational diffusion.

## Methods

### Overview

The diffusive and conformational behavior of a system of N beads simulated with the BD-HI algorithm of Ermak and McCammon [1] can be expressed in compact form as follows:

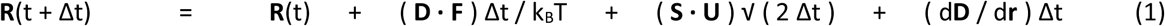

Here, **R**(t) is the 3N-dimensional vector storing the Cartesian coordinates of all N beads at time t, **D** is the 3N × 3N diffusion tensor describing the HIs between all bead pairs, **F** is the 3N vector storing the Cartesian components of the systematic forces acting on the beads, **S** is the 3N × 3N square root of the diffusion tensor (i.e. a matrix square root), or alternatively the diffusion tensor’s Cholesky decomposition, and **U** is a 3N vector of uncorrelated random displacements of zero mean and unit variance. The remaining non-trivial term, d**D** / d**r**, is a 3N vector that represents the divergence of the diffusion tensor (also termed the gradient of the tensor in the foundational Ermak and McCammon paper [1]); this term has the effect of translating each bead by an amount that represents the sum of all changes to the diffusion tensor due to changes in the position of all other beads in the system. Finally, Δt is the time step of the simulation, k_B_ is Boltzmann’s constant and T is the temperature in Kelvin.

The equations used to populate the elements of the diffusion tensor, **D**, are specified below, but first it is important to note where the computational expense of the BD-HI approach originates. The presence of the **D · F** term, which represents a matrix-vector product, indicates that the displacement of every bead is a function of the forces acting on every other bead in the system, with the diffusion tensor, **D**, determining the extent to which forces acting on one bead are converted into displacements of the other beads. Since **D** is a 3N × 3N matrix, the calculation of the force-induced displacements has a computational cost that scales, for large N, as (3N)^2^. A corresponding computational cost is incurred in the **S · U** term, which represents a second matrix-vector product that converts the 3N vector of uncorrelated random displacements **U** into a 3N vector of correlated random displacements, correlated in such a way as to ensure that the overall algorithm satisfies the fluctuation-dissipation theorem. In fact, the true computational cost associated with this second term is much greater than is suggested by a single matrix-vector product since we also need to include the computational cost of obtaining **S** from **D**; this operation scales as (3N)^3^, regardless of whether **S** represents the diffusion tensor’s matrix square root or its Cholesky decomposition. The expense of this operation is sufficiently great that for large systems this exact method is usually replaced by approximate, iterative methods that instead employ repeated matrix-vector products with a cost of (3N)^2^ (see below).

In order to illustrate the potential computational advantages of the method investigated here it is convenient to consider a simple system comprising only two beads of identical radius. For such a system, the calculation of the entire **D · F** term can be written as:

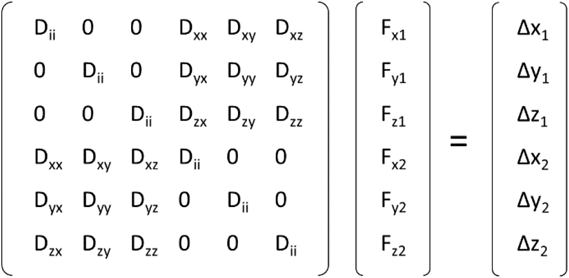

Here, the first bracketed term on the left is the 3N × 3N diffusion tensor, **D**, the second bracketed term is the 3N vector of force components, **F**, and the bracketed term on the right is the resulting 3N vector of force-induced displacements; for both of the 3N vectors, the subscripts 1 and 2 refer to beads 1 and 2, respectively. The D_ii_ terms represent the Stokes-Einstein translational diffusion coefficient describing the self-diffusion of each bead in each of the x,y,z directions (see below); these terms are determined by the radius of the bead and remain invariant throughout a BD-HI simulation. The D_xx_, D_xy_ etc terms represent the HIs as described by the RPY model for pairwise interactions of beads; these terms change throughout a BD simulation as the distance between the beads, and their relative orientations, change due to their motion. For larger systems, additional 3 × 3 submatrices would be included in **D** to describe the HIs between all pairs of beads.

The central idea explored here is to replace each 3 × 3 RPY tensor with its orientational average (OA), i.e. the average obtained by keeping the distance between the two beads fixed and sampling all possible angular orientations. This idea has been exploited by others previously in a variety of contexts (see Discussion), but to our knowledge, it has yet to be considered in the context of BD-HI simulations. As is made more explicit later, this orientational averaging has the effect of: (a) zeroing out all of the cross-terms in each 3 × 3 tensor, i.e. the D_xy_, D_xz_, etc terms, and (b) equating each of the remaining terms such that D_xx_ = D_yy_ = D_zz_ (= D_rr_; see below). For the two-bead system, the OA approximation simplifies the calculation of the **D · F** term to:

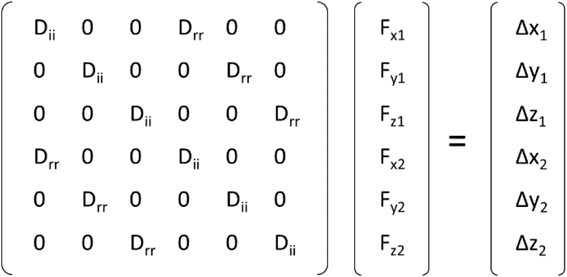

which in turn can be rearranged to:

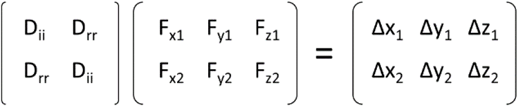

The total number of operations required to compute the **D · F** term with the complete (‘full’) RPY tensor is 3N × 3N multiplications and 3N additions, which becomes ∼9N^2^ when N is large. In comparison, the number of operations required with the OA version of the RPY tensor is N × N × 3 multiplications and 3N additions, which becomes ∼3N^2^ when N is large. All things being equal, therefore, we expect the OA RPY model to be three times faster than the full RPY model if the computer time required to perform each step of a BD simulation is dominated by the calculation of matrix-vector products.

While we immediately expect, therefore, that the OA RPY model will offer a computational advantage over the full RPY model for the calculation of the **D · F** term, its advantage with regard to the calculation of the correlated random displacements will likely depend on the system size. In equation (1), we wrote the calculation of the correlated random displacements as involving the product **S · U**, which assumes that the matrix square root or the Cholesky decomposition of **D** has been precomputed and is therefore available for use. This is conceptually the most straightforward way to obtain correlated random displacements: it works well for small and medium-sized systems, parallelized implementations of the Cholesky decomposition have become available over the years (e.g. [10]), and, in contrast to the faster, approximate treatments that are often used instead (see below), the correlated random displacements that are obtained reproduce exactly the statistical properties required for the fluctuation-dissipation theorem to be satisfied by the Ermak-McCammon algorithm. But the computational cost of obtaining a matrix square root or a Cholesky decomposition scales with the cube of the number of elements in each row (or column) of the matrix. For the full RPY model, then, this means that the cost scales as (3N)^3^, while for the OA RPY model, the cost scales instead as (N)^3^. For such situations, then, we can expect the Cholesky decomposition to run approximately 27 times faster under the OA approximation; reducing the practical effect of that speedup somewhat will be the fact that, in typical simulation studies, the expensive Cholesky decomposition is usually performed only at intervals of 5, 10, 100 or even 1000 simulation steps (e.g. [11]).

For larger systems, where the cubic scaling of the Choleksy decomposition becomes too expensive to tolerate, two alternative approximate methods are widely employed, both of which attempt to obtain progressively more accurate correlated random displacements by iteration. These are: (a) the Chebyshev polynomial method first proposed by Fixman [12], and lucidly described by the Graham and de Pablo groups [13], and (b) the Krylov subspace method developed and described by Chow, Skolnick and co-workers [11]; the latter method has the advantage that it does not require computation of the extreme eigenvalues of the diffusion tensor. While the details of these two methods differ – and the reader is referred to the original publications for information – the crucial point for the present work is that both involve repeated matrix-vector multiplications essentially identical in form to those examined earlier for the computation of the **D · F** term. As such, we can immediately expect the OA RPY model to have at least a 3-fold speed advantage over the full RPY model when either the Chebyshev or Krylov methods are employed. Additional data reported in Results indicates that this is likely to remain true in practical settings.

### Implementation details

In constructing the diffusion tensor **D** for a multi-bead system, the terms describing self-diffusion of beads are calculated using the standard Stokes-Einstein relationship:

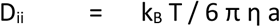

where a is the bead radius and η is the solvent viscosity. For the full RPY model, the equations specifying the 3 × 3 components of the diffusion tensor describing the HI of two beads of identical radius take the following forms:

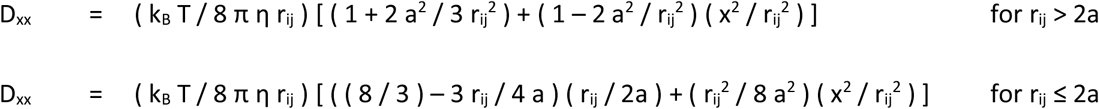

and:

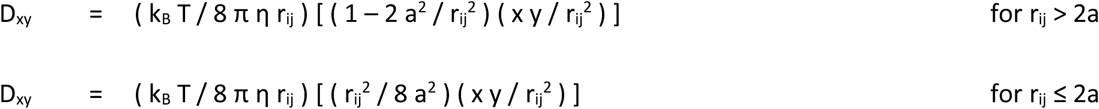

Here x, y, and z are the Cartesian components of the vector connecting the two beads, and r_ij_ is its magnitude. The remaining components of the diffusion tensor (D_xz_, D_yy_ etc) can be obtained by straightforward replacement of x, y and z in the above equations as appropriate. We note that while the present work exclusively considers situations in which all bead radii are identical, a comprehensive set of relations of RPY models for beads of unequal radii has been derived by the Szymczak group [14].

These equations simplify considerably when the components of the diffusion tensor are averaged over all possible orientations of the two beads, subject to the requirement that their distance remains constant. Specifically, terms of the form (x^2^ / r_ij_^2^) average to ⅓, while terms of the form (x y / r_ij_^2^) average to zero. As a result, regardless of the distance between the beads, D_xy_ = D_xz_ = D_yz_ = D_yx_ = D_zx_ = D_zy_ = 0, and the remaining three terms take the form:

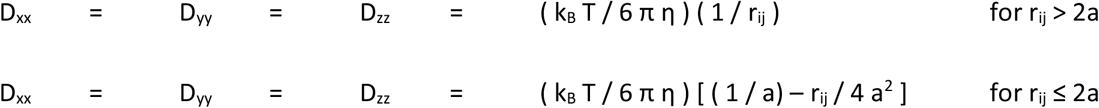

One important feature to note at this stage is that since the RPY model is known to produce a **D** that is positive definite (i.e. one that therefore can be subjected to Cholesky decomposition), the OA RPY model is also guaranteed to produce a positive definite **D**. The easiest way to see this is to consider the OA RPY **D** as the arithmetic average of an infinite number of **D**s, all calculated for the same system using the full RPY model but all differing in their overall orientation: a tensor that is the arithmetic average of tensors that are all positive definite is, by definition, also positive definite.

While the above equations are easy to implement, especially in simulation codes that already implement the full RPY tensor (see below), there is one final term to consider: this is the divergence term (d**D** / d**r**). With the full RPY model, this term is identically zero and so it is often omitted entirely from descriptions of the Ermak-McCammon algorithm. In the case of the OA RPY model, however, the divergence term is non-zero and takes the following form:

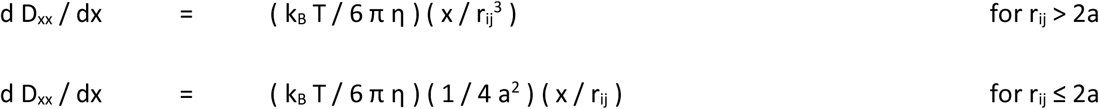

As above, displacement terms for the y and z coordinates use the same equations but with x replaced by y and z, respectively. The total displacement of each bead due to the (d**D** / d**r**) term is obtained by summing the contributions from all bead-bead interactions in the system. In principle, therefore, a BD-HI simulation performed with the OA RPY model incurs a minor additional cost that is not required in the full RPY model. The expected cost of this calculation is 3N^2^, but it is only performed when **D** is itself updated (see Results): at all other timesteps, the previously calculated divergence-related displacements are added unchanged.

### Simulations of residue-level coarse-grained models of proteins and RNAs

In order to test the ability of the OA RPY model to capture the translational and rotational diffusion of biological macromolecules, we selected eleven proteins and six RNAs that we have modeled in other studies (see Table S1). The proteins – which were previously modeled by us in a study exploring the use of the full RPY tensor in simulations of protein diffusion and folding [5] – range in size from 56 to 149 residues; the RNAs – which we have simulated in recent work describing a new method for building coarse-grained 3D models of very large RNAs [15] – range from 76 to 161 residues. For most of the simulations reported here we use residue-level coarse-grained representations: in the case of proteins, a bead is placed at the position of each Cα atom, while in the case of RNAs, a bead is placed at the position of each P atom. In all simulations that used residue-level representations, the form of the energy function used is essentially identical to that used in our previous works [5, 15, 16], which was itself closely modeled on that used by Clementi, Onuchic, and co-workers [17, 18]. Specifically, bonds are added between beads in adjacent residues, and standard molecular mechanics terms are then used to describe bond, angle, and dihedral angle deformations:

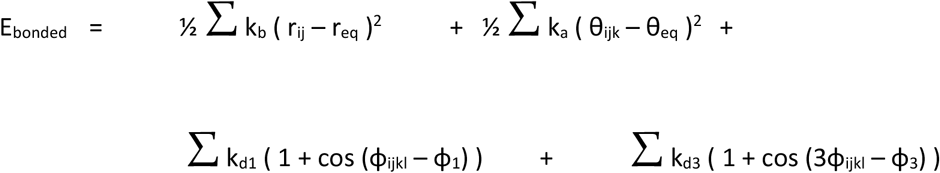

Here, the first summation is over all bonds in the molecule: r_ij_ is the current length of the bond between beads i and j, r_eq_ is the bond’s length at equilibrium, and k_b_ is the force constant, which was set to 20 kcal/mol/Å. The second summation is over all angles in the molecule: θ_ijk_ is the current angle between the i-j and j-k bond vectors, θ_eq_ is the angle’s value at equilibrium, and k_a_ is the force constant, which was set to 10 kcal/mol/rad. The third and fourth summations are over all dihedral angles in the molecule: the first of these two summations produces a cosine function with a single energy minimum and maximum; the second summation, which is added to add a degree of ‘roughness’ to the conformational energy landscapes [16–18], produces a cosine function with three energy minima and maxima, offset from each other by 120°. ϕ_1_ and ϕ_3_ are so-called “phase angles” that define the positions of the energy maxima. k_d1_ and k_d3_ represent the half-heights of the two dihedral energy functions; in all simulations, their values were set to 0.5 and 0.25 kcal/mol, respectively.

In addition to the above terms, the total interaction between all pairs of beads not involved in bonded interactions with each other was described using:

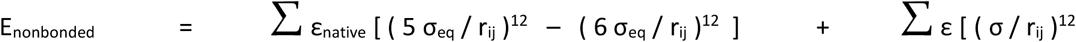

The first summation is a conventional Lennard-Jones “12-10” potential function evaluated for all pairs of beads that have at least one of their constituent atom pairs within 5.5 Å in the native state structure of the macromolecule; ε_native_ is the energy well depth, r_ij_ is the distance between the two beads, and σ_eq_ is the distance separating them in the native state structure. For simulations aimed at maintaining proteins and RNAs in their native (folded) states, ε_native_ was set to 1.0 kcal/mol; for simulations aimed at modeling their behavior while in unfolded states, ε_native_ was set to the much weaker value of 0.05 kcal/mol. The second summation is a conventional steric potential function that is applied to all nonbonded pairs of beads that are not considered to be in contact in the native state of the macromolecule; for simulations of proteins and RNAs, σ was set to 4 Å.

### Simulations of very coarse-grained models

Since it was of interest to determine whether the accuracy of the OA RPY model diminished as the level of resolution in the simulation models decreased, an additional set of simulations was performed for the largest protein considered here (SFVP) using a range of different resolutions. With the Cα model described above providing a resolution of 1 residue per bead, we made further lower-resolution models ranging from 3 residues per bead to 24 residues per bead. For each such model, beads were placed using the K-means approach implemented in the `qpdb` utility that is part of the Situs package [19]. Bonds were then added between the closest bead pairs until each bead had at least four such bonds; this was done to ensure that each model remained in a shape closely matching its initial structure throughout the simulations. Harmonic bond potential functions were then applied to each bond with a force constant of 20 kcal/mol/Å^2^; no angle or dihedral potential functions were applied in these simulations. The hydrodynamic radii of the beads were adjusted separately for each resolution model until their translational diffusion coefficients matched that of the Cα model when both were simulated with the full RPY model (see above).

### Brownian Dynamics simulations

All of the simulations reported here were performed using the *uiowa_bd* code written in-house (see below). Simulations of all proteins and RNAs were conducted for 10 µs using a timestep of 25 fs, and with coordinates saved for analysis at intervals of 100 ps. A nonbonded cutoff of 35 Å was used to construct a list of bead pairs within interacting distance, and this list was updated every 100 steps. Following our earlier work, the hydrodynamic radii of all protein beads in residue-level models were set to 5.3 Å as this value best reproduced the translational diffusion coefficient predicted for the same proteins using the hydrodynamics program HYDROPRO [20]. The hydrodynamic radii of all RNA beads were set to 5.5 Å as this value best reproduced the experimental translational diffusion coefficients of the same RNAs [15]. To explore the sensitivity of results to the frequency of updating the diffusion tensor, **D**, separate simulations were performed updating **D** at intervals of 25, 100, 250 and 1000 steps.

To measure the distribution of conformations sampled during the simulations, and to ensure correctness of the simulation algorithm (see Results), the radius of gyration, R_gyr_, of each conformation was measured using: R_gyr_^2^ = (1/N) Σ | r_i_ – r_mean_ |^2^, where the summation is over all N beads and r_mean_ is their mean position in that conformation. To examine the ability of the OA RPY model to reproduce dynamic properties predicted by the full RPY model, we chose to measure translational and rotational diffusion coefficients. To measure translational diffusion coefficients, D_trans_, the Einstein formula was used: < r^2^ > = 6 D_trans_ δt, where < r^2^ > is the mean-squared distance traveled by the molecule’s center of geometry during an observation interval δt, which was set here to 10 ns [21]. Rotational diffusion coefficients were obtained as follows. First, each folded state CG model was aligned along its principal axes of inertia using the GROMACS [22, 23] utility `princ`. Next, pairs of beads were identified whose inter-bead vector best matched the x-, y-, and z principal axes, respectively. Such bead pairs were identified by examining all possible bead-pairs, retaining only those pairs whose separation distance along the axis of interest was at least 75% of the maximum separation distance along the same axis found for any pairs, and selecting the pair with the smallest combined distance from the axis of interest. With representative bead pairs identified for each of the principal axes, rotation of each axis during the simulations was monitored by calculating the autocorrelation function: θ(t) = < **e**_**i**_(t) · **e**_**i**_(0) >, where **e**_**i**_ is the relevant normalized principal axis vector at time t [24]. For the proteins, a single exponential function of form y = exp (– δt / τ_rot_), was fit to each correlation function up to δt = 10 ns, and the final rotational diffusion coefficient for rotation of that axis was obtained as D_rot_ = 1 / 2 τ_rot_. For the RNAs, an identical approach was used but owing to the slower rotation, the fit to each correlation function was carried out up to δt = 56 ns. Error estimates for D_trans_ and D_rot_ values were obtained by calculating each quantity separately for three equal-sized blocks of each trajectory: 0.1 – 3.4, 3.4 – 6.7, and 6.7 – 10.0 µs, and then obtaining the standard deviation of the three values.

## Code availability

All BD simulations described here were performed using the in-house code *uiowa_bd*, earlier versions of which have been used in a number of simulation studies [5, 16, 25–32]. The source code for *uiowa_bd* is available at the following GitHub repository (https://github.com/Elcock-Lab/uiowa_bd).

## Results

Since there are good theoretical reasons to expect that the OA RPY model will be considerably faster to compute than the full RPY model (see Methods), the purpose of the remainder of this manuscript is to assess the extent to which the model provides a description of macromolecular diffusion that is sufficiently close to that obtained from the full RPY approach to warrant using in future simulations of diffusion and folding events. Following our previous work, and reflecting our own application areas of interest, we focus here on modeling the translational and rotational diffusion of folded and unfolded proteins and RNAs, each represented as bead-spring models in which each bead represents one amino acid or nucleotide, respectively. Throughout this manuscript we use the full RPY model as our “gold standard”, since it is the more complete RPY theory that we intend the OA RPY approximation to mimic.

### The OA RPY Model Provides A Reasonable Description Of HIs

Before examining the impact of the OA RPY approximation on the simulated behavior of macromolecules, it is instructive to compare the nature of the force-induced displacements that are predicted by the OA RPY model with those predicted by the full RPY model. One way to do this is to consider the system shown in Figure 1. Here, we imagine a hypothetical 2D arrangement of beads, with the blue spheres representing the beads’ initial positions, and the red spheres representing their final positions after the **D · F** term has been applied. We consider a situation in which only the central bead (circled) experiences any systematic force, in this case a force in the x-direction that is sufficient to move it as shown; the “bonds” connecting the blue and red spheres show how each of the beads will be displaced. The panel on the left shows the behavior obtained when the **D · F** term is calculated using the full RPY model for each HI; the panel on the right shows the behavior obtained when the **D · F** term is instead calculated using the OA RPY model. A number of similarities and differences between the two models are apparent. With the full RPY tensor, the force-induced displacements of the beads surrounding the central bead behave in line with intuition: (a) beads are either pushed away from, or dragged along with, the central bead, (b) beads with a diagonal relationship to the central bead are themselves displaced diagonally, and (c) the magnitude of the displacements decreases with increasing distance from the central bead. With the OA RPY model, the first and last of these observations are reproduced, but, as expected from a model that involves orientational averaging, the diagonal displacements are lost: instead, the displacements of all other beads are co-directional with that of the central bead, and with magnitudes determined entirely by the distance of each bead from the central bead. Comparison of the two figure panels immediately suggests, therefore, that the OA RPY model might perform somewhat better at reproducing the full RPY model’s translational diffusion than rotational diffusion; this idea is borne out below.

**Figure 1.**
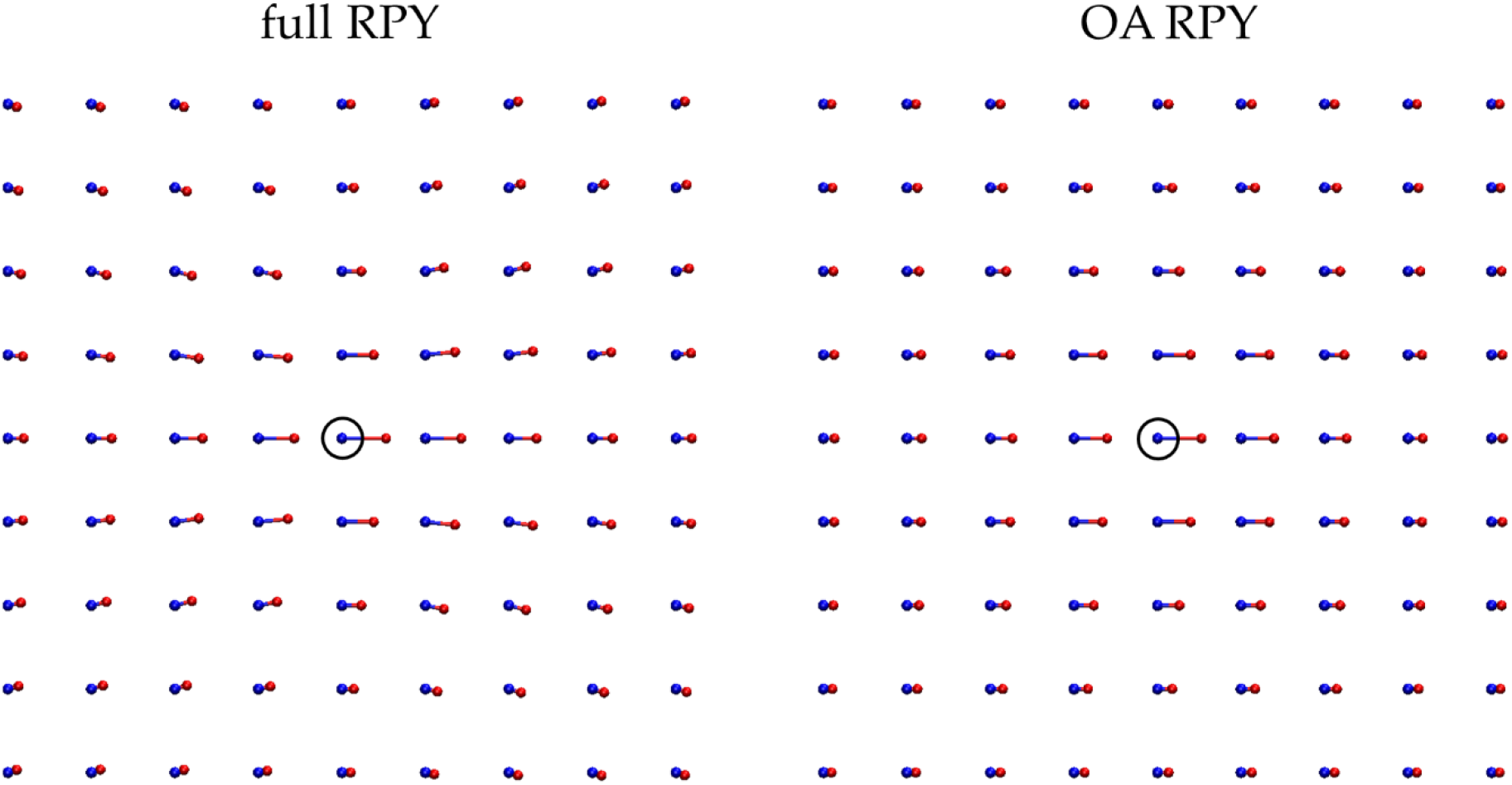
Comparison of the force-induced displacements obtained from the full RPY model (left) with those obtained from the OA RPY model (right). Spheres show the positions before (blue) and after (red) application of the **D · F** term calculated when the only force acting in the system is one acting on the central bead (circled). For these calculations, all beads were assigned a hydrodynamic radius of 5.3 Å and were separated from their nearest neighbor by 5.3 Å.

### OA RPY Matches Full RPY Only When the Divergence Term is Included

In order to determine the correctness of our implementation of the OA-RPY model in BD-HI simulations, we considered first the static properties of eleven proteins modeled in their unfolded states (Figure 2a). As noted in Methods, on a purely theoretical basis, the use of the OA RPY model requires the inclusion of a non-zero divergence term that, in conventional applications of the full RPY model would otherwise be identically zero. Here we show that the divergence term is not only required in principle, but also absolutely required in practice if reasonable results are to be obtained. Figure 2b shows scatterplots comparing the mean radii of gyration for the eleven proteins simulated for 10 µs with energetic parameters that ensure they sample unfolded conformations. When the divergence term is included (blue symbols), the mean R_gyr_ values obtained with the OA RPY model match exactly with those obtained with the full RPY model (in this and all other Figures, the diagonal line corresponds to y=x). Just as importantly, the very wide range of conformations sampled during the simulations – which is reflected in the large error bars which represent the standard deviation of the sampled R_gyr_ values – is the same in both the OA RPY and full RPY simulations. In contrast, when the divergence term is omitted (red symbols), the mean R_gyr_ values obtained with the OA RPY model are far too small, indicating that the supposedly unfolded proteins have collapsed unrealistically into highly compact states. We conclude, therefore, that the divergence term is an essential component to include in simulations that aim to use the OA RPY model. Interestingly, this is the case even though the magnitude of displacements caused by the divergence term are themselves very small: in simulations of the largest protein, SFVP, for example, the mean and maximum magnitudes of the divergence term’s displacement on the first step of a simulation of the protein in its native state were only 0.00175 and 0.00206 Å, respectively.

**Figure 2.**
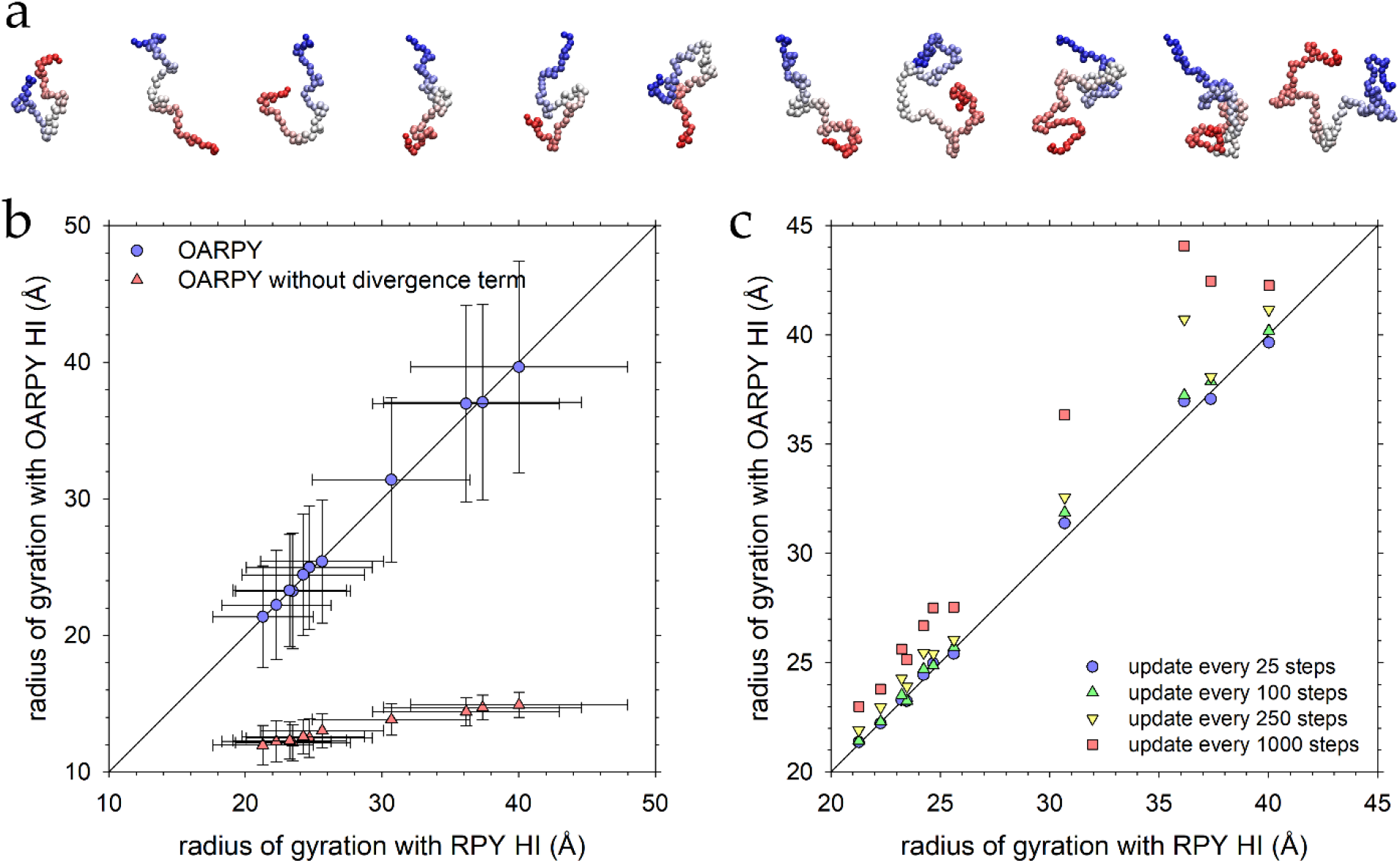
(a) Snapshots of the eleven unfolded proteins taken at the end of a 10 µs BD-HI simulation. (b) Comparison of the mean radius of gyration values (R_gyr_) obtained from simulations of unfolded proteins performed with the OA RPY model with those obtained with the full RPY model. In all simulations, the diffusion tensor and divergence terms were updated every 25 simulation steps. Data are shown for simulations performed with (blue) and without (red) the divergence term required by the Ermak-McCammon algorithm (see main text). Error bars represent the standard deviation of the full distribution of the R_gyr_ values calculated for all 9900 conformations obtained for each protein. The solid line represents y = x. (c) Comparison of the mean radius of gyration values obtained from simulations of unfolded proteins performed with the OA RPY model (including the divergence term) with those obtained with the full RPY model. For the OA RPY model, data are shown for four different update intervals for the diffusion tensor and the divergence terms; for the full RPY model, the update interval was fixed at 25 steps.

We next sought to address how often the divergence term must be updated during simulations. In typical applications of the full RPY model in BD-HI simulations it is common to keep the diffusion tensor unchanged for some number of simulation steps (e.g. 10 to 100 simulation steps): this is especially common in applications where the Cholesky decomposition is used to calculate correlated random displacements (see Methods). When the RPY model is used, updating **D** infrequently should have little or no impact on static properties such as the R_gyr_ distribution (although it may affect dynamic properties such as rotational diffusion coefficients). We explicitly demonstrate this in Figure S1 where we show that the mean R_gyr_ values obtained from simulations with the RPY model are effectively unchanged regardless of whether **D** is updated at intervals of every 25, 100, 250 or 1000 steps. But with the OA RPY model it might be anticipated that more frequent updating might be required since the divergence terms are constant displacements repeatedly added to the beads at every simulation step. This indeed appears to be the case. Figure 2c shows that the ability of the OA RPY model to match the mean R_gyr_ values obtained with the full RPY model diminishes as the number of simulation steps between each update of **D** (and the divergence terms) increases. When the update frequency is 25 steps, the absolute deviation of the mean R_gyr_ values obtained with the OA RPY model from those of the full RPY model average to only 0.98%; when the update frequency increases to 100 steps, the error rises to 1.33%, when the update frequency rises to 250 steps, the error reaches 4.13%, and when the update frequency rises to 1000 steps (i.e. every 25 ps) the error reaches 11.0%. Because of these results, we assumed for all of the following simulations that **D** should be updated at intervals of 100 simulation steps.

### Translational Diffusion Of Proteins Is Better Described By The OA RPY Model Than Rotational Diffusion

Having established that static properties of unfolded proteins can be reproduced successfully by our implementation of the OA RPY model, we then sought to determine how well the model reproduced the dynamic properties predicted by the full RPY model for proteins modeled in both their unfolded states (Figure 2a) and their folded states (Figure 3a). Figure 3b shows scatterplots comparing computed D_trans_ values obtained using the OA RPY model with those obtained using the full RPY model for proteins modeled in their unfolded states; Figure 3c does the same for the proteins in their folded states. The agreement is excellent in both cases, and this impression is amplified when the ratio of the folded to unfolded D_trans_ values is considered. In our previous work [5] we showed that HIs are essential to include in BD simulations if they are to reproduce the experimental observation that proteins diffuse ∼30-60% faster in their folded states than in their unfolded states; Figure 3d demonstrates that the OA RPY model also accurately reproduces this behavior. Finally, Figure 3e compares the rotational diffusion coefficients obtained with the OA RPY model to those obtained with the full RPY model; note that separate values are computed for rotation of each of the 3 principal axis vectors in each protein (see Methods). Qualitatively, the agreement is excellent, but quantitatively it is clearly worse for rotational diffusion than for translational diffusion: the rotational diffusion coefficients predicted by the OA RPY model are ∼25% lower than they should be, with mean absolute percent errors in the D_rot_ values of 26.4, 23.4, and 24.3 % for rotation of the x-, y-, and z-axis vectors, respectively. Interestingly, the data plotted in Figure 3e hint that these errors might increase in relative terms as the “true” (i.e. full RPY) D_rot_ value increases, since the datapoints appear to curve away from the y=x line. But this suggestion is not strongly supported by further analysis of the data: when the 33 absolute percent errors are plotted versus the corresponding RPY D_rot_ value, the r^2^ of the linear regression is only 0.063 (Figure S2a). Nor is there any obvious dependence of these errors on the size of the protein: a linear regression of the mean absolute percent errors versus the number of beads in the protein models produces an r^2^ value of only 0.023 (Figure S2b).

**Figure 3.**
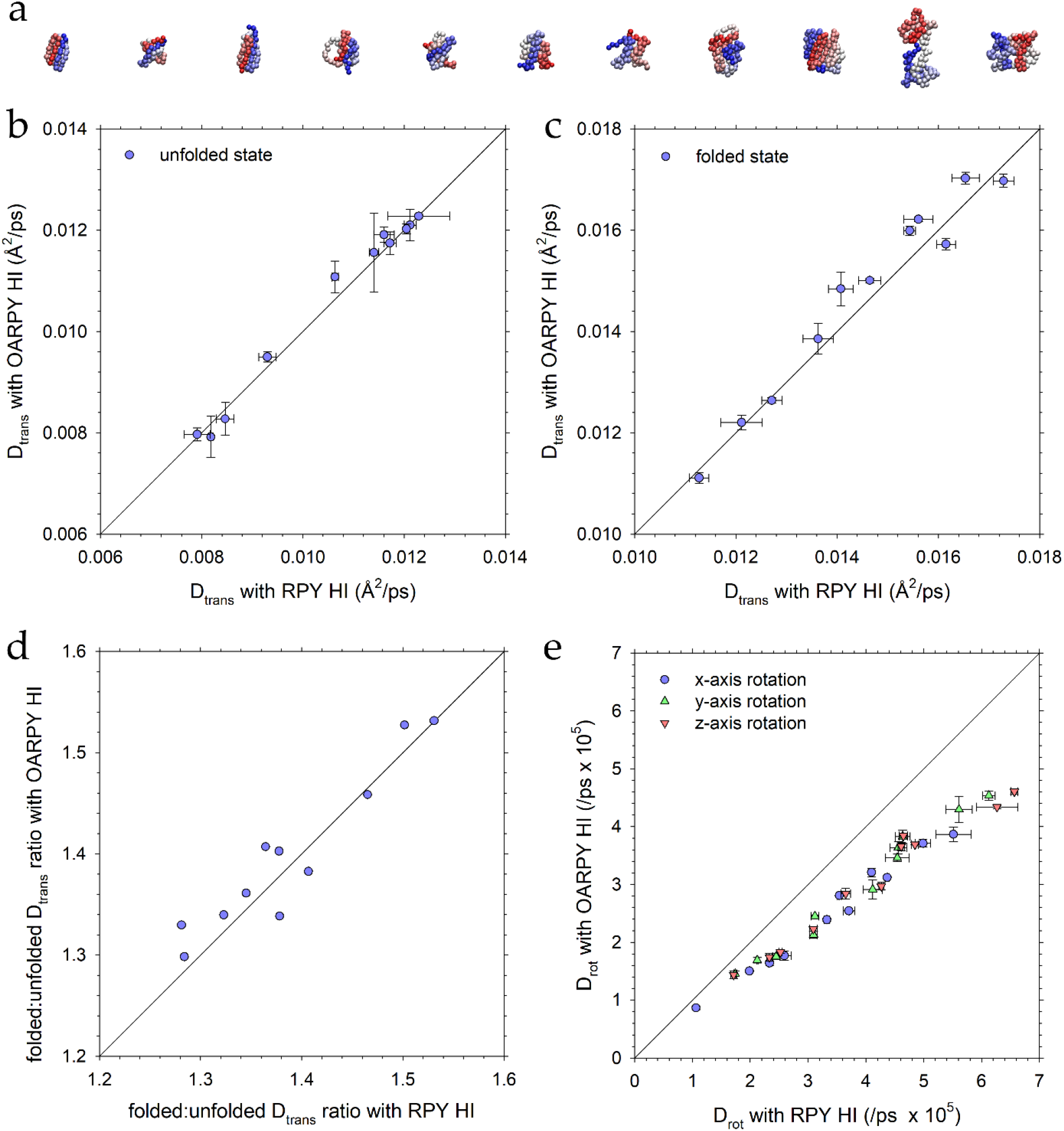
(a) Structures of the eleven folded proteins. (b) Comparison of the translational diffusion coefficient (D_trans_) obtained from simulations of unfolded proteins performed with the OA RPY model with those obtained with the full RPY model. In this and all subsequent figures, error bars are calculated as described in the main text. (c) Same as (b) but showing results for the same proteins simulated in their folded states. (d) Ratio of D_trans_ measured in the folded and unfolded states for the OA RPY model plotted versus the same ratio obtained with the full RPY model. (e) Comparison of the rotational diffusion coefficient (D_rot_) for each of the principal axes obtained from simulations of folded proteins performed with the OA RPY model with those obtained with the full RPY model.

### The Accuracy Of OA-RPY With RNAs Is Similar To That With Proteins

The results reported thus far provide a strong indication that the OA RPY model can accurately approximate a protein’s diffusive characteristics, at least in comparison with the full RPY model. While there is no compelling basis for imagining that qualitatively different results would be obtained with different kinds of macromolecules, we were interested in determining whether the errors in rotational diffusion predicted by the OA RPY method would be magnified in molecules whose shape is more asymmetric than the predominantly spherical nature of most of the globular proteins in our dataset. To explore this issue, we conducted additional simulations using six different RNA molecules (Figure 4a), five of whose translational diffusion coefficients we have recently shown [15] match their experimental values very closely when simulated with the full RPY model. The additional RNA simulated here is an 85 nucleotide stem loop that is ∼130 Å long, with a diameter of ∼18 Å; we add this in order to test the extent to which the OA RPY model might work with very elongated molecules. Scatterplots comparing the translational and rotational diffusion coefficients obtained using the OA RPY model with those obtained using the full RPY model are shown in Figure 4b and 4c, respectively. The results are qualitatively identical to those seen with the proteins: the D_trans_ values are again reproduced very well, while the D_rot_ values are consistently underestimated, in this case by 20%, relative to the full RPY values. We conclude that the OA RPY model remains equally valid for simulations of RNAs as it does for proteins.

**Figure 4.**
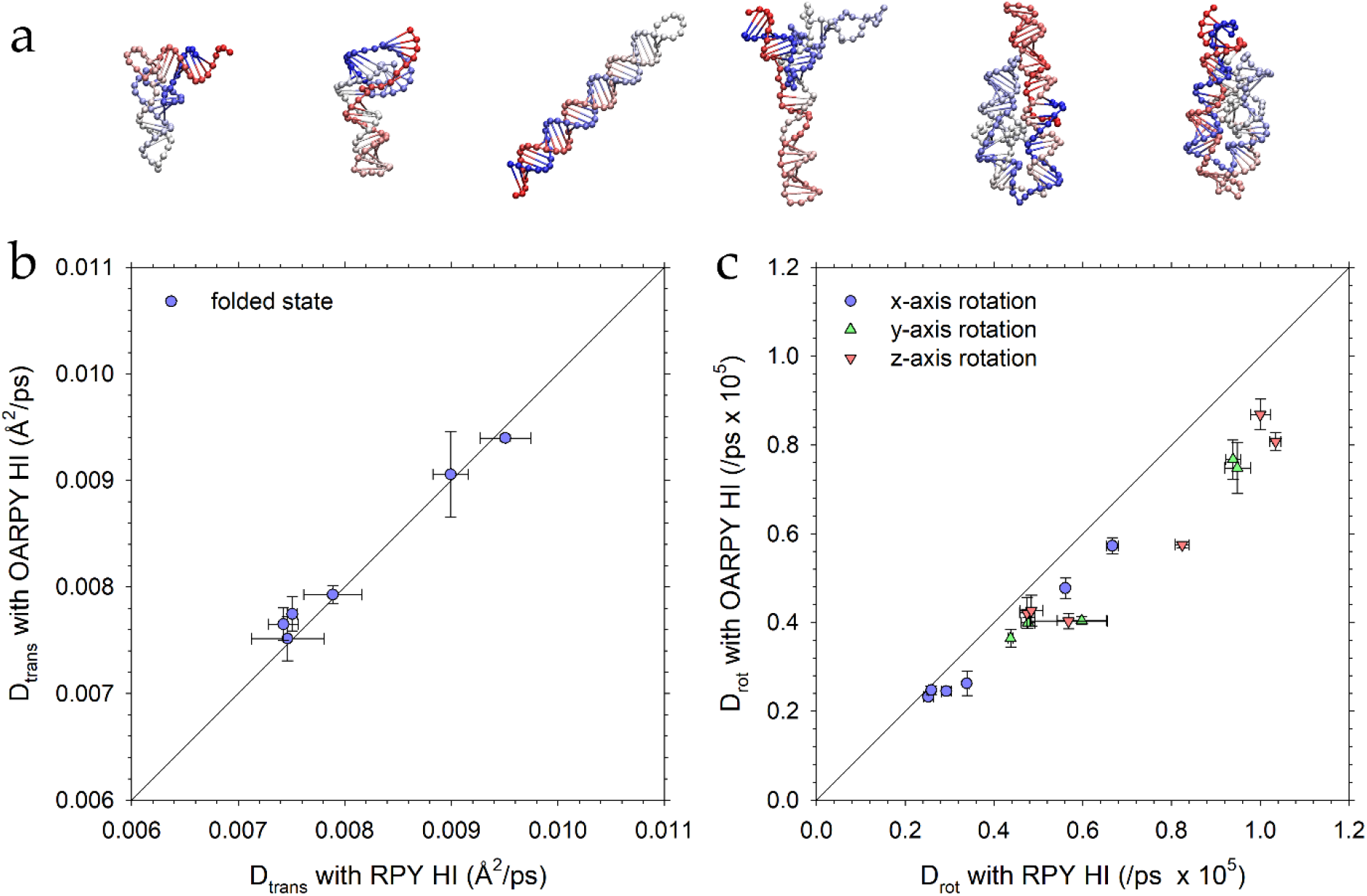
(a) Structures of the six folded RNAs. (b) Comparison of the translational diffusion coefficient (D_trans_) obtained from simulations of folded RNAs performed with the OA RPY model with those obtained with the full RPY model. (c) Comparison of the rotational diffusion coefficient (D_rot_) for each of the principal axes obtained from simulations of folded RNAs performed with the OA RPY model with those obtained with the full RPY model.

### The Accuracy of the Rotational Diffusion Coefficient is Independent of Resolution

The results presented thus far have indicated that the OA RPY model works well for both proteins and RNAs when they are modeled at resolutions of 1 bead per amino acid and 1 bead per nucleotide, respectively. Since one possible future use of the OA RPY model, however, could be to larger systems that might contain many copies of much more coarsely-modeled macromolecules, it was of interest to determine whether the model’s ability to match the full RPY model might diminish as the resolution of the models decreased. To explore this issue, we selected the largest of the proteins in our dataset, the 149-residue protein SFVP, and made 9 different models with progressively lower levels of resolution starting at one-bead-per-residue and proceeding all the way down to one-bead-per-24-residues, which contains a total of only 6 CG beads (Figure 5a). For each resolution, we first adjusted the bead radius in the full RPY simulations to roughly match the translational diffusion coefficient obtained with our highest-resolution model: this ensures that all the different resolution models are normalized according to their translation. We then used the same bead radius in a simulation performed with the OA RPY model. Figure 5b compares the D_trans_ values obtained for each of the 9 CG models of SFVP as simulated with both the OA RPY and full RPY models; Figure 5c does the same for the D_rot_ values of the x-axis (plots for the y- and z-axes, which tell the same story, are shown in Figures S3a and S3b). Regardless of the resolution of the CG model, the same trends are obtained as before, i.e. the D_trans_ values predicted by the OA RPY model match very closely with those predicted by the corresponding full RPY model, while the D_rot_ values are underestimated by ∼20%. We conclude, therefore, that the OA RPY model is likely to be a viable, fast alternative to the full RPY model for a wide range of CG representations of macromolecules.

**Figure 5.**
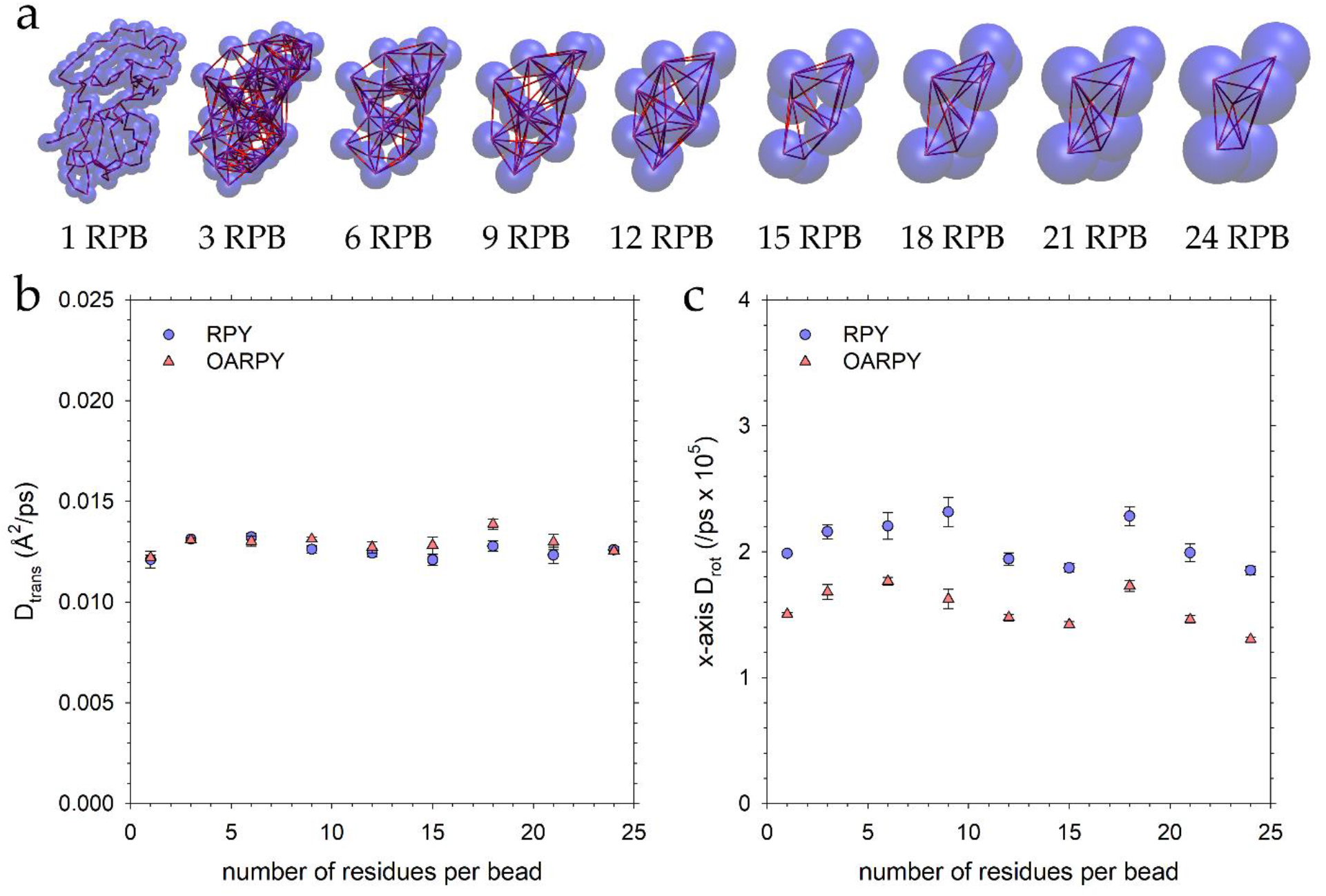
(a) Structures of the nine different models of SFVP that were simulated. The resolutions range from: (left) 1 residue per bead (1 RPB) to (right) 24 residues per bead (24 RPB). (b) Plot of the translational diffusion coefficient (D_trans_) versus the resolution of the coarse-grained model for (blue) the full RPY model and (red) the OA RPY model. For each resolution, the bead radius for the full RPY model was first adjusted in an attempt to match the D_trans_ value obtained with 1 RPB; the same bead radius was then used in the corresponding simulation performed with the OA RPY model. (c) Same as (b) but showing the rotational diffusion coefficient (D_rot_) of the principal x-axis of the molecule. The same bead radii were used to obtain the results in (b) and (c).

### A Note on the Use of the OA-RPY model with the Chebyshev polynomial method

As noted in the Introduction, the computational benefits of using the OA RPY model are likely to be greatest for intermediate-scale systems in which the Cholesky decomposition remains the most efficient means of (exactly) computing correlated random displacements. For larger-scale systems, where the (approximate) Chebyshev and/or Krylov-based methods become the method of choice, the computational gains of the OA RPY model are expected to be reduced from ∼27-fold to ∼3-fold. But the latter number assumes that the computational cost of all other aspects of the Chebyshev and/or Krylov-based methods remain unchanged. For the Chebyshev method, in particular, it is well known that the rate at which converged estimates of the random displacements are obtained is a function of the ‘condition number’ of the diffusion tensor, i.e. the ratio of the diffusion tensor’s maximum eigenvalue to that of its minimum eigenvalue: as the condition number increases, so the number of iterations required for convergence also increases [11, 13]. To be certain, therefore, that the OA RPY model is likely to retain its 3-fold competitive advantage over the full RPY model, we should make sure that the condition number of the OA RPY diffusion tensor is less than or equal that of the full RPY diffusion tensor. To explore this issue, we computed the condition numbers for folded and unfolded conformations of each of the 11 proteins simulated here, using both the OA RPY and full RPY models. Importantly, the condition numbers of the OA RPY diffusion tensors are all somewhat lower than those of the full RPY diffusion tensors (see Figure 6). While these differences are probably not so large that they would give the OA RPY model any additional advantage over its theoretical 3-fold speedup in using the Chebyshev method, they do indicate that the 3-fold speedup is likely to be secure. Furthermore, while the Krylov method does not formally require the extreme eigenvalues to be computed, its rate of convergence is likely also to exhibit some sensitivity to the condition number.

**Figure 6.**
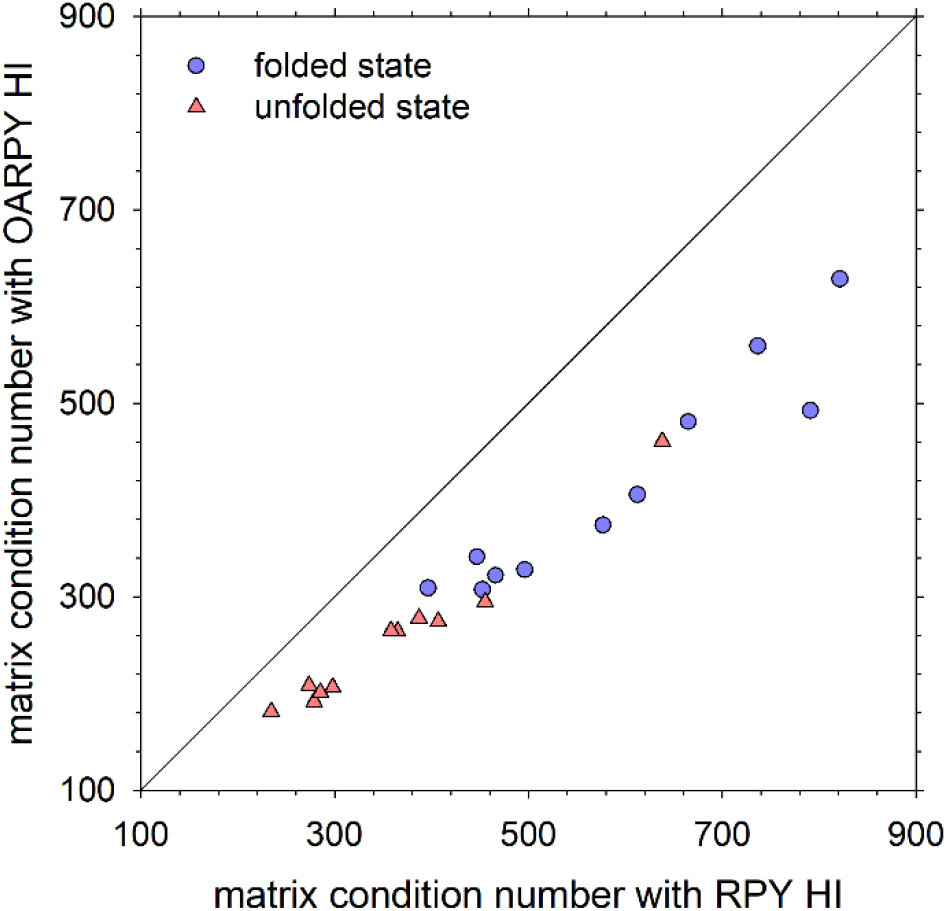
Comparison of the condition number of the diffusion tensor, **D**, for the eleven proteins using the OA RPY model with that obtained using the full RPY model. Datapoints are shown for proteins in their folded states (blue circles) and unfolded states (red triangles).

## Discussion

Use of the full RPY HI model in combination with various kinds of coarse-grained molecular representations is a well-established method to simulate the diffusion and conformational dynamics of biomolecules [4, 5, 27, 33–35], their folding kinetics (e.g. [36–38]), and their association rate constants (e.g. [28, 39]). In Methods, we made the case that the OA RPY model has clear computational advantages that make it worth investigating as a means of accelerating BD-HI simulations. In Results, we have shown that these advantages come with few attendant disadvantages: simulations of isolated proteins and RNAs indicate that translational diffusion coefficients are very accurately reproduced while rotational diffusion coefficients are consistently underestimated by ∼25%. Depending on the system under study, it may be that the latter drawback is a price worth paying for the increased speed provided by the OA approximation.

While the current work is, to our knowledge, the first to explore the use of an OA RPY tensor in BD-HI simulations, this is certainly not the first time that orientational averaging of HIs has been considered in the literature. For example, orientational averaging of the Oseen tensor was a key step in the derivation of Kirkwood and Riseman’s 1948 theory of the intrinsic viscosity and translational diffusion of flexible polymers [6]. It is also a key component of studies that connect the equations of classical electrostatics with those of hydrodynamics and exploit the former to more easily calculate transport coefficients of rigid body models of macromolecules (e.g. [8, 40, 41]). Interestingly, rigid body calculations – not based on the use of BD simulations – by the groups of Garcia de la Torre [9] and Aragon [8] have both already shown that orientational averaging allows translational diffusion coefficients to be accurately reproduced but causes a significant underestimation of rotational diffusion coefficients (or equivalently, an overestimation of rotational friction coefficients). Clearly, those results prefigure a number of the results obtained here.

The clearest connections with the current work, however, are to be found in: (a) the Sing group’s use of an iterative conformational averaging approach that averages the HIs not only over orientations but also over distances [42], and (b) the Garcia de la Torre group’s study of the effects of orientational averaging on the diffusional properties of flexible polymers [7]. The interesting method developed by the Sing group has the advantage, once converged, of never requiring an update to either **D** or **S** but this comes at the expense of not allowing fluctuations in the magnitudes of the HIs. The latter aspect makes the approach less appropriate for use in BD-HI simulations that attempt to model the folding behavior of proteins given that their shapes change drastically upon folding. In the work of the Garcia de la Torre group, Monte Carlo methods were used to generate many polymer conformations (an approach originally pioneered by Zimm [43]) and each was then treated as a rigid body in order calculate translational and rotational friction coefficients and obtain ensemble averages. Again, translational friction was found to be unaffected by invoking the OA approximation while rotational friction was found to be significantly overestimated. Interestingly, for a freely-jointed chain polymer model, the extent of the error in rotational friction was dependent on the ratio of the beads’ hydrodynamic radius (σ) to the average bond length (b), with the results getting progressively worse as this ratio increased. In the protein simulations described here, σ is 5.3 Å and b is 3.8 Å, giving a ratio, σ/b, of 1.39. This is much higher than the highest ratio considered in the calculations of Garcia de la Torre *et al*. (0.3922), but if we use a logarithmic equation to regress their reported data for the largest freely jointed chain in their data set and extrapolate to a σ/b ratio 1.39, we obtain an estimated percent error in the rotational friction of 19%, which is similar to the error in rotational diffusion found here in BD-HI simulations (Figure 3e).

For large biomolecules, the idea of using Monte Carlo or other fast methods to generate possible conformations and then carrying out hydrodynamic calculations under the assumption that each behaves like a rigid body is likely to continue to be useful for estimating transport properties, particularly in cases where performing BD-HI simulations becomes very expensive [34]. But given BD-HI simulations’ other uses, especially in explicitly simulating folding or association events, the development of methods to accelerate BD-HI simulations remains an important goal. We have shown here that a key technical issue that arises when applying the OA RPY in BD-HI simulations is the need to include a term related to the divergence of the diffusion tensor. This term is often omitted from discussion in BD-HI studies since it is zero when the full RPY tensor is used [1]. But several studies have already demonstrated the importance of this term in situations different from the current one. For example, Grassia, Hinch and Nitsche showed how inclusion of the Ermak-McCammon divergence term (referred to there as the “mean drift” term) is needed to obtain correct behavior in conditions where the diffusivity of a particle varies with its position [44]. Heyes, in developing an innovative mean-field treatment of hydrodynamics intended to accelerate the simulation of concentrated colloidal systems, also noted the need for including a friction-related term that is equivalent to the divergence term used here [45]. Finally, the Donev group has shown that a corresponding “stochastic drift” term, which they show can be calculated using a random finite difference method, is also necessary for correct Boltzmann sampling in BD-HI simulations of confined particle suspensions [46, 47]. Given these prior works, the results obtained here showing that the divergence term is required for the OA RPY model to produce correct static properties (Figure 2b) is not a surprise. Fortunately, the term is easily calculated, and so long as it is updated at sufficiently frequent intervals (Figure 2c), it can be added unchanged with no obvious error in simulated behavior.

One additional result that we have obtained here is that the OA RPY model’s ability to describe translational and rotational diffusion is essentially unaffected by the level of coarse-graining in the structural models. It is perhaps worth emphasizing that this result is true also for the full RPY model: bead radii that are optimized to reproduce the translational diffusion coefficient predicted by a much finer-resolution model also allow very coarse-grained models to reproduce the finer model’s rotational diffusion, even when the coarse model contains as few as six beads (Figure 5c). That the OA RPY model’s relative ability to reproduce rotational diffusion does not deteriorate further as the resolution of the model decreases suggests that it is likely to be useful even in large-scale applications where very coarse-grained structural models might be used.

In closing, it is worth considering the potential applications of the OA RPY model. For reasons outlined earlier, the exact speedup to be obtained will likely depend on the situation, so the question of whether the speedup is worth the price of a ∼25% poorer reproduction of rotational diffusion will have to be answered on a case-by-case basis. One potential use that can certainly be suggested, however, is during the “equilibration” period of large-scale BD-HI simulations. For very large systems, such as complex mixtures of macromolecules whose initial positions might have been assigned randomly (e.g. [48, 49]), the time period required to lose memory of the initial configuration might be very substantial. Since we have explicitly shown that static properties obtained with the OA RPY model match exactly those obtained with the full RPY model, the OA RPY model could be used to accelerate this part of the simulation without any penalty. If more accurate reproduction of rotational diffusion is ultimately desired, the full RPY model could then be substituted during the “production” phase of such simulations.

## Figure generation

All molecular images were generated with VMD [50]. All graphs were generated with SigmaPlot (Systat Software, San Jose, CA).

## Acknowledgments

This research was supported by a grant from the National Institutes of Health (R35 GM122466) to AHE and supported in part through computational resources provided by The University of Iowa.

## Author Contributions (CRediT)

John W. Tworek: Formal analysis, Investigation, Writing – original draft preparation, Visualization.

Adrian H. Elcock: Conceptualization, Methodology, Software, Formal analysis, Investigation, Writing – review and editing, Supervision, Project Administration, Funding acquisition.

